# Area-specific mapping of binocular disparity across mouse visual cortex

**DOI:** 10.1101/591412

**Authors:** Alessandro La Chioma, Tobias Bonhoeffer, Mark Hübener

## Abstract

Binocular disparity, the difference between left and right eye images, is a powerful cue for depth perception. Many neurons in the visual cortex of higher mammals are sensitive to binocular disparity, with distinct disparity tuning properties across primary and higher visual areas. Mouse primary visual cortex (V1) has been shown to contain disparity-tuned neurons, but it is unknown how these signals are processed beyond V1. We find that disparity signals are prominent in higher areas of mouse visual cortex. Preferred disparities markedly differ among visual areas, with area RL encoding visual stimuli very close to the mouse. Moreover, disparity preference is systematically related to visual field elevation, such that neurons with receptive fields in the lower visual field are overall tuned to near disparities, likely reflecting an adaptation to natural image statistics. Our results reveal ecologically relevant areal specializations for binocular disparity processing across mouse visual cortex.

## Introduction

Depth perception is a fundamental feature of many visual systems across species. It is relevant for many behaviors, like spatial orientation, prey capture, and predator detection. Multiple cues contribute to depth perception, among them binocular disparity, the small difference between left and right eye images, arising when a stimulus is viewed by two eyes. Binocular disparity changes as a function of object distance from the observer and can hence provide the visual system with critical information for depth perception (Gonzalez and Perez 1998; Cumming and DeAngelis 2001). In primates, individual neurons sensitive to binocular disparity are found throughout most of the visual cortex, with different disparity tuning properties across primary and higher visual areas, suggesting specific roles of different higher areas for depth perception (Gonzalez and Perez 1998; Cumming and DeAngelis 2001; Parker 2007).

Mouse primary visual cortex (V1) has been shown to contain disparity-tuned neurons, similar to those found in other mammals (Scholl et al. 2013; 2015), but it is unknown how binocular disparity is processed beyond V1 and whether it is differentially represented in higher areas. Mouse visual cortex comprises V1 and more than a dozen higher-order areas that show partially different tuning properties for basic stimulus features, like spatial and temporal frequency and motion direction (Andermann et al. 2011; Marshel et al. 2011; Roth et al. 2012; Glickfeld and Olsen 2017; Murakami et al. 2017; Smith et al. 2017). However, the specific roles of these higher areas for visual information processing are not well understood. Apart from V1, areas LM and RL contain the largest representation of the binocular visual field (Garrett et al. 2014; Zhuang et al. 2017), making them candidate areas for investigating downstream processing of binocular disparity in mouse visual cortex. In turn, comparison of disparity tuning across different mouse visual areas might help delineating their functional specializations.

## Results

To investigate binocular disparity tuning in mouse visual cortex, areas V1, LM, and RL were identified and localized using intrinsic signal imaging (Marshel et al. 2011; Garrett et al. 2014). By using the established visual field representations in mouse visual cortex, the boundaries between V1, LM, and RL could be readily identified (Fig. S1; see Methods for details). We then targeted the binocular regions of areas V1, LM, and RL for functional two-photon imaging (Fig. S1), and used the genetically encoded calcium indicator GCaMP6s to measure visually-evoked activity of individual neurons (Chen et al. 2013).

### Disparity sensitivity is widespread across areas of mouse visual cortex

Disparity tuning was initially characterized using drifting vertical gratings displayed in a dichoptic fashion at varying interocular disparities (Fig. 1A; Scholl et al. 2013). Twelve different grating disparities were generated by systematically varying the relative phase of the two gratings presented to either eye, while drift direction, speed, and spatial frequency were kept constant across eyes. Across areas V1, LM, and RL, about one third of neurons were responsive (see Methods) to vertical, dichoptic gratings, with comparable response magnitudes across areas (percentage of responsive cells, mean ± SEM across imaging planes, V1: 34.6% ± 1.2%, LM: 32.3% ± 1.9%, RL: 34.2% ± 1.5%; ΔF/F in response to gratings, mean ± SEM across planes, V1: 53% ± 2%, LM: 43% ± 2%, RL: 48% ± 2%; number of imaging planes, V1: n=52; LM: n=18; RL: n=33).

**Figure 1.**
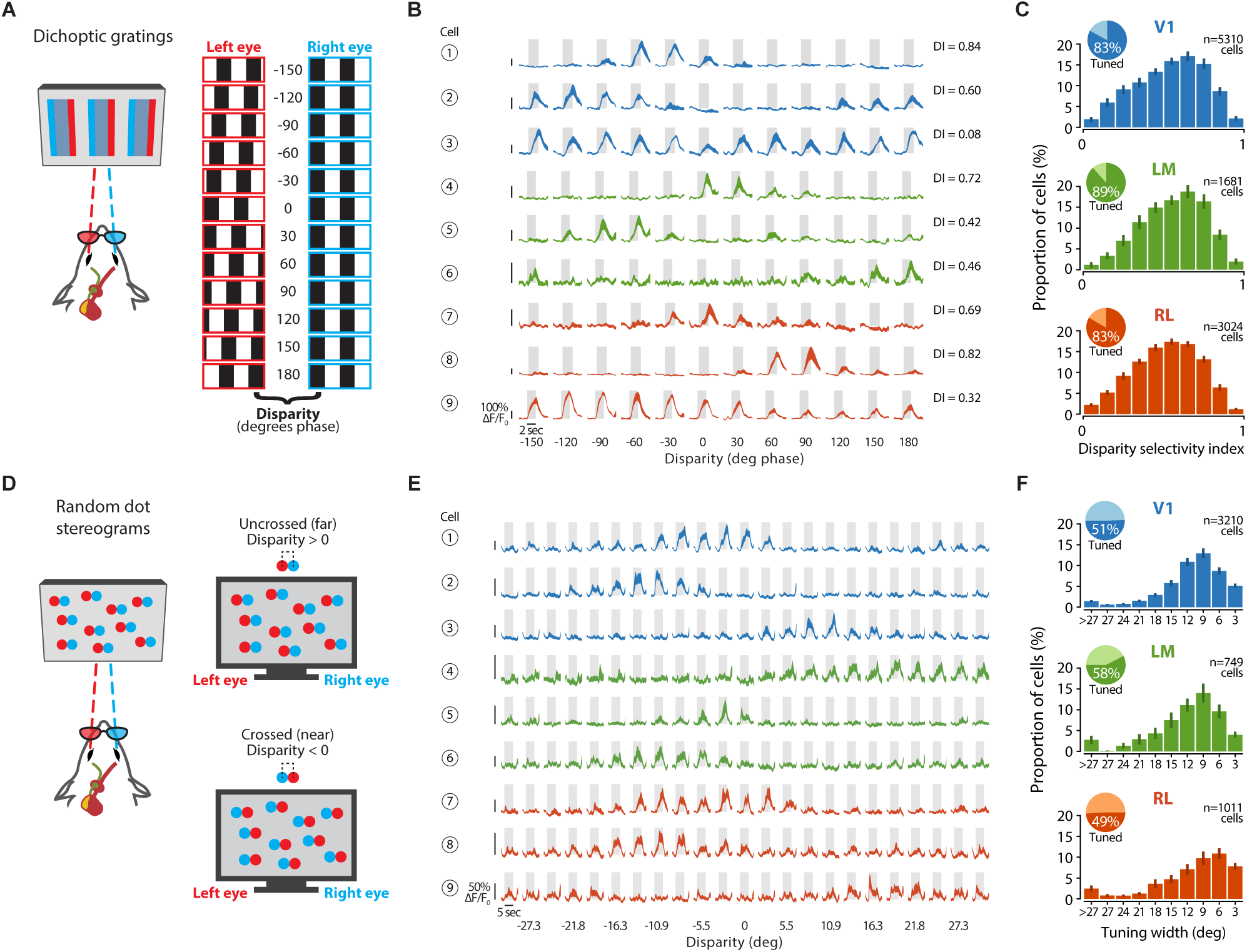
Disparity sensitivity is widespread across areas of mouse visual cortex. (A) Schematic of drifting grating stimuli, presented dichoptically through eye shutter glasses. Twelve equally spaced interocular grating disparities (−150–180 deg phase, spacing 30 deg phase) are generated by systematically varying the initial phase between the gratings presented to either eye. The red-cyan color code illustrates the eye-specific presentation of the gratings; in reality, gratings are displayed in black and white. (B) Visually-evoked calcium traces (ΔF/F_0_) of nine example neurons from the three areas (here and throughout: blue, V1; green, LM; orange, RL). For each neuron, the disparity selectivity index (DI) is indicated on the right. Fluorescence time courses are plotted as mean ΔF/F_0_ ± SEM (shaded areas) calculated across stimulus trials (5-6 repeats). Gray boxes, duration of stimulus presentation (2 sec), bottom edge indicates baseline level (0% ΔF/F_0_). (C) Distributions of disparity selectivity index (DI) for each area. Mean of medians across planes ± SEM, V1: 0.55 ± 0.01, LM: 0.58 ± 0.02, RL: 0.53 ± 0.01; Kruskal-Wallis test, *χ*^2^=3.870, p=0.144 (V1: n=52 planes, 105.1 ± 4.1 cells per plane, 5310 responsive cells total, 16 mice; LM: n=18 planes, 93.4 ± 5.0 cells per plane, 1681 responsive cells total, 7 mice; RL: n=33 planes, 91.6 ± 4.7 cells per plane, 3024 responsive cells total, 13 mice). Inset, percentage of disparity tuned and untuned cells (see Methods); percentage tuned cells mean ± SEM across planes, V1: 83.1% ± 1.5%, LM: 88.5% ± 1.5%, RL: 83.4% ± 1.6%; Kruskal-Wallis test, *χ*^2^=3.855, p=0.146. (D) Random dot stereogram (RDS) stimuli. Left, schematic of the dichoptic presentation of RDS stimuli through eye shutter glasses. Top right, positive horizontal displacements cause binocular disparities equivalent to far object distances. Bottom right, negative horizontal displacements cause binocular disparities equivalent to near object distances. (E) Example calcium traces (9-10 repeats) in response to RDS stimuli of nine different neurons from areas V1, LM and RL. (F) Distributions of disparity tuning width for each area, determined with RDS as the width parameter of the Gaussian fit (see Methods). Mean of medians across planes ± SEM, V1: 8.61 deg ± 0.29 deg, LM: 9.19 deg ± 0.88 deg, RL: 7.65 deg ± 0.65 deg; Kruskal-Wallis test, *χ*^2^= 4.004, p=0.135 (V1: n=38 planes, 84.5 ± 6.3 cells per plane, 3210 responsive cells total, 16 mice; LM: n=13 planes, 57.6 ± 9.2 cells per plane, 749 responsive cells total, 6 mice; RL: n=19 planes, 53.2 ± 5.8 cells per plane, 1011 responsive cells total, 11 mice). Inset, percentage of tuned and untuned cells; percentage tuned cells mean ± SEM across planes, V1: 50.6% ± 3.1%, LM: 57.6% ± 4.5%, RL: 49.1% ± 3.8%; Kruskal-Wallis test, *χ*^2^=2.086, p=0.352.

For each responsive cell, a disparity tuning curve was computed by plotting its average calcium response as a function of the interocular phase disparity of the grating (Fig. 1B). Across areas, many neurons showed a strong modulation of disparity tuning. To quantify the magnitude of disparity tuning, a disparity selectivity index (DI) was calculated for each cell, with values closer to one for highly selective cells and values closer to zero for unselective cells (Fig. 1B). Cells were defined as disparity-tuned in response to gratings when their DI values were above 0.3. To determine whether any of the three areas was specifically dedicated for encoding binocular disparity, we plotted the frequency distribution of DI values for each area (Fig. 1C). While in each area the majority of neurons showed at least some degree of disparity tuning (>80% disparity-tuned neurons), the overall degree of disparity selectivity was similar across areas.

Next we probed disparity tuning in mouse visual cortex using random dot stereograms (RDS; Fig. 1D; Julesz 1971). RDS have been instrumental for investigating disparity sensitivity in primates, but have never been employed in mice. Unlike gratings, RDS allow measuring absolute disparities, in which tuning curves are expressed as a function of visual angle, avoiding the ambiguity deriving from the circular nature of gratings. In addition, RDS are spatially more homogeneous than gratings, lacking any motion and orientation signals, and thus allowing a better isolation of the disparity component of a cell’s response.

RDS stimuli generally activated neurons less strongly than drifting gratings (percentage of responsive cells, mean ± SEM across planes, V1: 28.6% ± 2.1%, LM: 20.1% ± 3.2%, RL: 19.5% ± 2.2%; ΔF/F in response to RDS, mean ± SEM across planes, V1: 44% ± 1%, LM: 37% ± 2%, RL: 43% ± 2%; number of imaging planes, V1: n=38; LM: n=13; RL: n=19). Still, many neurons exhibited clear responses to RDS, with reliable activation by a limited range of disparities (Fig. 1E). Approximately half of the neurons that responded to RDS were disparity-tuned (see Methods), showing clear disparity selectivity, with comparable fractions of cells and tuning sharpness across areas (Fig. 1F). Thus, binocular disparity signals are prominent not only in mouse V1 (Scholl et al. 2013; 2015), but also in higher areas LM and RL.

### Area RL is specialized for encoding near disparities

While disparity selectivity was overall similar across areas, we found clear differences in disparity preference measured with dichoptic gratings: area RL contained a significantly higher fraction of neurons tuned to negative (near) disparities compared to areas V1 and LM (Fig. 2A). The over-representation of negative disparities in area RL compared to V1 and LM was also evident by plotting the averaged disparity preference for each imaging plane, showing a consistent difference across animals and imaging sessions (Fig. 2B). Note that a crossed disparity of e.g. –30 deg phase is equivalent to an uncrossed disparity of 330 deg phase, owing to the correspondence problem created by dichoptically presented, phase-shifted gratings. Despite this ambiguity, at a spatial frequency of 0.01 cycles per degree, as used here, neurons are more likely activated by the smaller of the two possible phase disparities (–30 deg rather than 330 deg), given the typical receptive field (RF) size in mouse visual areas (Van den Bergh et al. 2010; Smith et al. 2017) and a binocular overlap of about 40 deg (Scholl et al. 2013). Thus, the data obtained with dichoptic gratings suggest that area RL encodes disparities corresponding to visual stimuli closer to the mouse, compared to V1 and LM.

**Figure 2.**
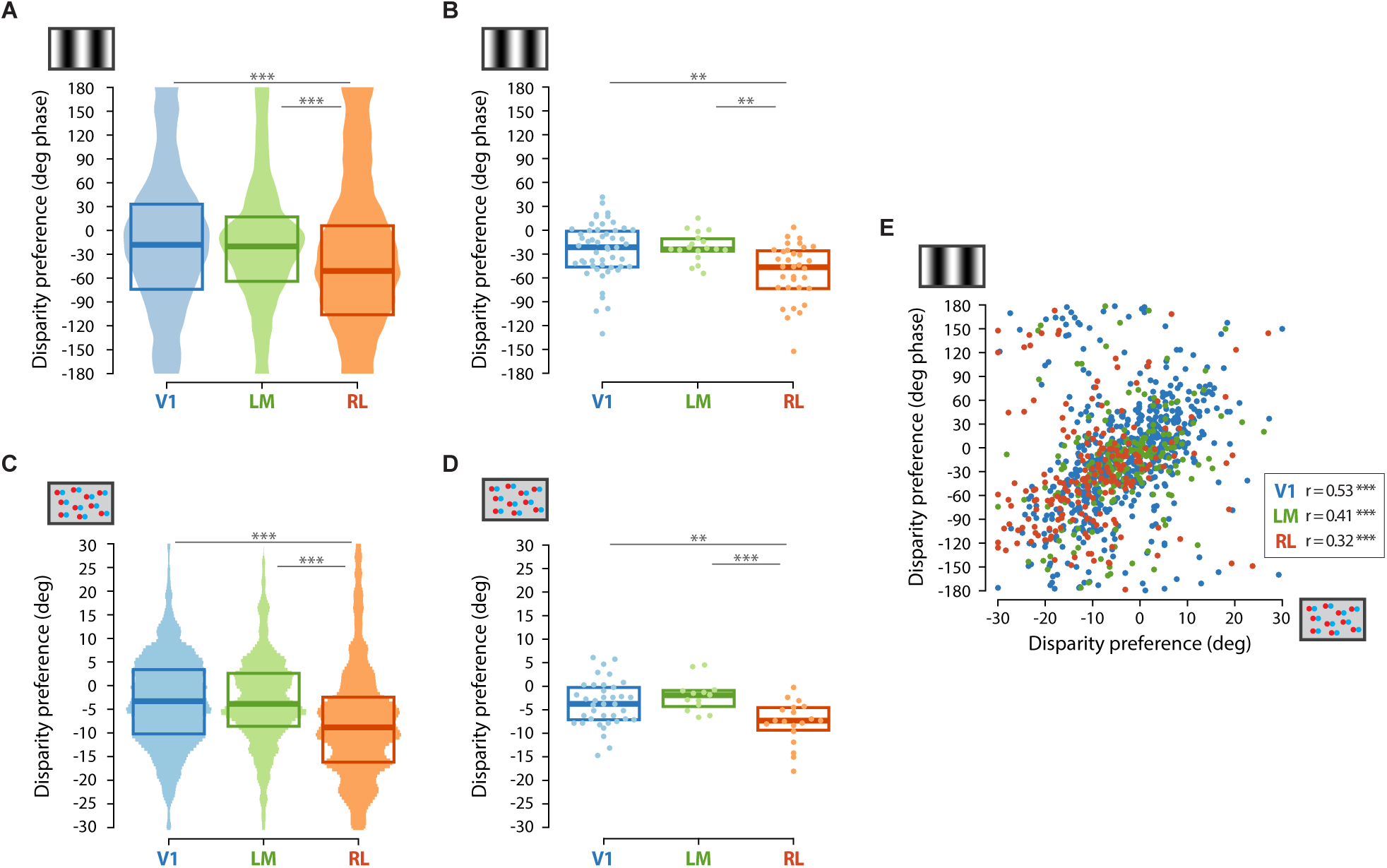
Area RL is specialized for encoding near disparities. (A) Disparity preference measured with gratings, across all disparity-tuned neurons in each area. Box plots indicate median and interquartile range calculated across neurons. Shaded areas show the mirrored circular probability density estimates (kernel width 10 deg) normalized to the number of data points in each area (V1: n=4070 cells, 52 planes, 16 mice; LM: n=1348 cells, 18 planes, 7 mice; RL: n=2326 cells, 33 planes, 13 mice; Watson-Williams test, F(2,7741)=92.98, p<1e-16; Bonferroni-corrected post hoc Watson-Williams tests: V1 vs. LM, p=0.4650, V1 vs. RL, p<1e-15, LM vs. RL, p<1e-15). (B) Disparity preference measured with gratings, data points are averages across neurons in each imaging plane. Box plots indicate median and interquartile range calculated across imaging planes. V1: n=52 planes, 16 mice; LM: n=18 planes, 7 mice; RL: n=33 planes, 13 mice; Watson-Williams test, F(2,100)=8.12, p=5.4050e-04; Bonferroni-corrected post hoc Watson-Williams tests: V1 vs. LM, unadjusted p=0.7437, V1 vs. RL, p=2.4086e-03, LM vs. RL, p=3.8166e-03). (C) Disparity preference measured with RDS, across all disparity-tuned neurons in each area. Box plots indicate median and interquartile range calculated across neurons. Shaded areas show the mirrored probability density estimates (kernel width 3 deg) normalized to the number of data points in each area (V1: n=1645 cells; LM: n=454 cells; RL: n=511 cells; Kruskal-Wallis test: *χ*^2^=99.377, p=2.6341e-22; Bonferroni-corrected post hoc tests: V1 vs. LM, unadjusted p=0.9934, V1 vs. RL, p=8.0583e-22, LM vs. RL, p=7.6514e-14). (D) Disparity preference measured with RDS, data points are averages across neurons in each imaging plane. Box plots indicate median and interquartile range calculated across imaging planes. V1: n=38 planes; LM: n=13 planes; RL: n=19 planes; Kruskal-Wallis test: *χ*^2^=13.239, p=0.0013339; Bonferroni-corrected post hoc tests: V1 vs. LM, p=0.5525, V1 vs. RL, p=0.0130, LM vs. RL, p=0.0019. (E) Correlation between disparity preference determined with gratings and RDS, for individual neurons tuned to both stimuli. Correlation coefficient between a circular and a linear variable, V1: r=0.531, p<1e-15; LM: r=0.409, p=3.8682e-08; RL: r=0.3187, p=3.4980e-05 (V1: n=644 cells, 39 planes; LM: n=204 cells, 17 planes; RL: n=202 cells, 25 planes).

To directly prove this interpretation, we next used RDS to measure absolute disparity preferences in the three areas. Indeed, we found that neurons in area RL were significantly shifted towards negative (near) values compared to cells in V1 and LM (Fig. 2C,D). Tuning for negative disparities in RL was also seen in awake animals using both grating and RDS stimuli (Fig. S2). Thus, neuronal signals recorded in response to two very different visual stimuli, dichoptic gratings and RDS, show that area RL is specialized for encoding binocular disparities corresponding to nearby visual stimuli.

Finally, we tested whether disparity preferences determined with dichoptic gratings or RDS are correlated at the level of individual neurons. For neurons tuned to both stimuli, disparity preference was highly correlated in all three areas (Fig. 2E). Thus, both types of stimuli are well suited to assess neurons’ disparity tuning, and both stimuli demonstrate distinct preferences for binocular disparity across visual areas.

### Disparity preference is related to visual field elevation

Although the binocular regions of areas V1, LM, and RL contain similar representations of the visual field in azimuth, they cover partially different regions of the visual field in elevation (Garrett et al. 2014; Zhuang et al. 2017). While binocular V1 represents both the lower and the upper visual field, the retinotopic representations of LM and RL cover mainly the upper and lower visual field, respectively. Thus, the distinct preference of RL for near disparities, as compared to V1 and LM, might be related to the different visual field representations that these areas show in elevation, rather than arising from dedicated, area-specific processing.

To directly assess the potential relationship between disparity tuning and RF elevation, multiple planes (2 to 4) within the same imaging session were recorded in the binocular regions of V1 and RL, aligned along the rostrocaudal axis and hence at distinct retinotopic elevations (Fig. 3A). For individual neurons, we determined RF elevation using horizontal bars of drifting gratings displayed at various vertical locations (Fig. 3B,C). For the same neurons, we also measured disparity tuning, using either dichoptic gratings or RDS, as described above.

**Figure 3.**
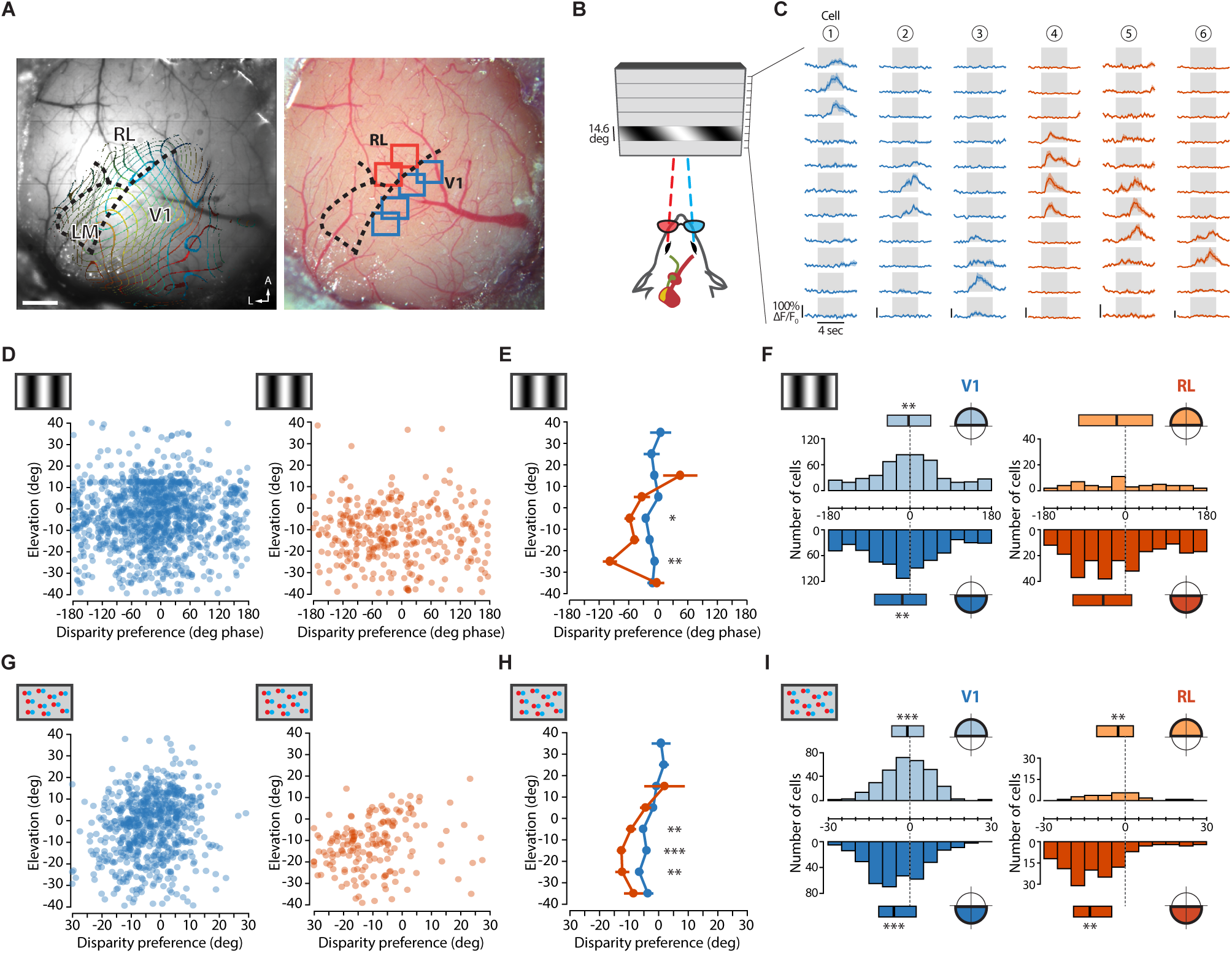
Disparity preference is related to visual field elevation. (A) Image of the cortical surface acquired through a cranial window. Left, contour plots of azimuth and elevation overlaid on an image of the brain surface. Boundaries between areas V1, LM, and RL are indicated with dashed black lines. Right, locations of imaging planes in V1 (blue, four planes) and RL (orange, two planes). Scale bar, 500 µm. (B) Schematic of horizontal bars of drifting gratings for measuring RF position in elevation. Individual bars have a vertical size of 14.6 deg and are presented in steps of 7.3 deg. (C) Example calcium traces (ΔF/F_0_) of six neurons in response to horizontal bar stimuli as shown in (b). (D) Correlation between disparity preference measured with gratings and RF position in elevation, for individual neurons in V1 (blue) and RL (orange). Correlation coefficient between a circular and a linear variable, V1: r=0.046, p=0.282; RL: r=0.021, p=0.933 (V1: n=1200 cells, 27 planes; RL: n=317 cells, 8 planes). (E) Disparity preference measured with gratings averaged across neurons for each elevation bin (bin width, 10 deg). Permutation tests (see Methods): [−40 −30] deg, p=0.778; [−30 −20] deg, p=0.0009; [−20 −10] deg, p=0.080; [−10 0] deg, p=0.0343; [0 10] deg, p=0.143; [10 20] deg, p=0.291. (F) Distributions of disparity preference measured with gratings plotted separately for upper and lower visual field. Median and interquartile range of disparity preferences for upper and lower visual field are also shown (medians, V1 up: −3 deg phase; V1 low: −17 deg phase; RL up: −19 deg; RL low: −49 deg; Watson-Williams test, V1 up vs. V1 low, F(1,1198)=7.89, p=0.0051; RL up vs. RL low, F(1,315)=2.30, p=0.1301). Vertical dashed lines indicate zero disparity. (G) Correlation between disparity preference measured with RDS and RF position in elevation, for individual neurons in areas V1 (blue) and RL (orange). Spearman correlation coefficient, V1: r=0.194, p=4.2274e-07, n=672 cells, 20 planes; RL: r=0.236, p=1.8463e-03, n=172 cells, 8 planes. (H) Disparity preference measured with RDS averaged across neurons of each bin of RF elevation. Permutation tests (see Methods): [−40 −30] deg, p = 0.3814; [−30 −20] deg, p = 0.0075; [−20 −10] deg, p < 1e-04; [−10 0] deg, p = 0.0060; [0 10] deg, p = 0.2600; [10 20] deg, p = 0.5287. (I) Distributions of disparity preferences measured with RDS plotted separately for upper and lower visual field. Median and interquartile range of disparity preferences for upper and lower visual field are also shown (medians, V1 up: −0.9 deg; V1 low: −5.9 deg; RL up: −2.6 deg; RL low: −13.1 deg; Kolmogorov-Smirnov test, V1 up vs. V1 low: D=0.2313, p=2.2906e-08; RL up vs. RL low: D=0.3919, p=9.7125e-04). Vertical dashed lines indicate zero disparity.

For individual disparity-tuned cells in areas V1 and RL, there was no linear correlation between disparity preference as measured with gratings and RF position in elevation (Fig. 3D). Nonetheless, V1 cells with RFs located in the lower half of the visual field showed a disparity preference that was significantly more negative than V1 cells with RFs in the upper visual field (p=0.0051), pointing to a relation between disparity tuning and retinotopic elevation (Fig. 3E,F). Neurons in area RL showed a similar, albeit not significant (p=0.1301), relationship: cells with RFs in the lower visual field were on average tuned to more negative disparities than cells with RFs in the upper visual field (Fig. 3E,F). Note that relatively few cells in RL had their RFs in the upper visual field, consistent with RL covering mostly the lower visual field. When using RDS for stimulation, a systematic variation of disparity preference with retinotopic elevation became evident for both areas V1 and RL (Fig. 3G). Accordingly, neurons in both V1 and RL with RFs in the lower visual field were significantly (p<0.001) tuned to more negative disparities compared to cells with RFs located in the upper visual field (Fig. 3H,I). Importantly, across corresponding locations in the lower visual field, neurons in RL were on average tuned to significantly more negative (near) disparities compared to cells in V1 (Fig. 3E,H). Thus, the functional specialization of RL for encoding near disparities is not simply related to its retinotopic representation of the lower visual field, but is likely mediated by area-specific disparity processing.

## Discussion

Our data show that binocular integration endows many neurons in the binocular regions of areas V1, LM, and RL of mouse visual cortex with sensitivity for retinal disparities. The respective representations of binocular disparities exhibit systematic differences, with area RL being specialized for close visual stimuli, compared to V1 and LM.

The disparity preferences observed using RDS allow estimating the absolute distances of visual objects encoded by the three areas. Considering the stereo-geometry of the mouse visual system (Fig. S3A,B; Scholl et al. 2013), and using the interquartile ranges of disparity preference values as depicted in Fig. 2C, area V1 encodes visual objects at distances between 3.6 and 24.9 cm from the mouse’s eyes, area LM between 4.0 and 16.7 cm, and area RL between 2.6 and 6.8 cm (Fig. S3C). Thus, disparity-tuned neurons in area RL are tuned to a narrow range of distances, very close to the mouse. This range of distances is well within reach of the mouse’s whiskers, which can be as long as ~3 cm (Ibrahim and Wright 1975; Brecht et al. 1997), originating from the whisker pad that is located ~1 cm distal to the eyes. In fact, mouse RL, located between V1 and S1 barrel cortex, is not a unimodal visual, but rather a multisensory area that harbors layer- and cell-specific circuits mediating visuo-tactile integration (Olcese et al. 2013) and receives strong, direct projections from V1 and the barrel cortex (Gămănuţ et al. 2018).

While the topographic organization of whisker driven inputs to area RL is not known, the upper layers of the barrel cortex itself contain a highly ordered, continuous map of near touch-space, which becomes overt during active sensation with all whiskers intact (Pluta et al. 2017). Given the prominent whisker driven inputs to area RL (Olcese et al. 2013; Gămănuţ et al. 2018), it is plausible that also neurons in area RL are organized into an orderly map for touch-space. Since (lower) visual space is mapped systematically across RL (Garrett et al. 2014; Zhuang et al. 2017) (Fig. S3), with a preference for near objects, as shown here, we speculate that area RL contains matched vision and touch sensory maps to form a multimodal representation of near space in front of the mouse, potentially important for object interaction, jointly mediated by vision and active whisking. In primates, vision and arm/hand movements are used in a coordinated fashion for object interaction, reflected by activity of neurons in the “parietal reach region” within the posterior parietal cortex (Rizzolatti et al. 1990; Andersen and Buneo 2002). Of note, area RL in the mouse has been proposed to be part of the rodent posterior parietal cortex (Hovde et al. 2018).

In both areas, V1 and RL, we found a clear relationship between disparity preference and retinotopic elevation: neurons with RFs in the upper visual field are tuned to more positive (far) disparities, while neurons with RFs in the lower visual field are driven by more negative (near) disparities (Fig. 3). While the potential correlation between disparity tuning and retinotopic elevation has not been directly assessed in cats or monkeys, a recent study compiling data from several publications in macaque V1 (Sprague et al. 2015), and a recent fMRI study in humans (Nasr and Tootell 2018) found the same relationship that we report here for the mouse. Not surprisingly, the natural distribution of binocular disparities measured in humans exhibits a clear gradient in elevation, especially during object interaction, with negative (near) disparities prevalent in the lower and positive (far) disparities in the upper visual field (Sprague et al. 2015). While not yet directly measured, it is obvious that a similar distribution of disparities characterizes natural viewing in the mouse, too, thereby matching the elevation-dependent distribution of disparity preferences as reported here. Thus, as it has been shown for other visual RF properties (Geisler 2008), also binocular response properties of cortical neurons tightly reflect the statistics of natural images, suggesting that visual experience sculpts RF properties during development for achieving the best adaptation to a three-dimensional visual environment.

## Acknowledgements

We thank Pieter Goltstein for setting up intrinsic signal imaging hard- and software and for comments on an earlier version of the manuscript. We are grateful to Max Sperling for the development of data acquisition software. This work was supported by the Max Planck Society and a Boehringer Ingelheim Ph.D. fellowship to A.L.C.

## Author contributions

A.L.C. and M.H. designed the project. A.L.C performed the experiments and analyzed the data. All authors wrote the manuscript.

## Declaration of interests

The authors declare no competing interests.

## Methods

### Ethics

All experimental procedures were carried out in accordance with the institutional guidelines of the Max Planck Society and the local government (Regierung von Oberbayern).

### Virus injection and cranial window implantation

Cranial window implantations were performed on 16 female adult C57/BL6 mice (10–13 weeks of age at the start of the experiment). Mice were anesthetized by intraperitoneal injection of a mixture of Fentanyl (0.075 mg/kg), Midazolam (7.5 mg/kg), and Medetomidine (0.75 mg/kg). A general analgesic (Carprofen, 4 mg/kg, subcutaneous injection) was administered immediately before surgery and for two days during post-surgical recovery. After an initial skin incision, a local analgesic (Lidocaine 10%) was topically applied. To facilitate the positioning of the craniotomy and enhance the accuracy and reproducibility of the targeted virus injections, intrinsic optical imaging was performed through the skull to coarsely locate the binocular region of V1. A circular craniotomy (4-5 mm diameter) was performed with a dentist drill, centered over the binocular region of V1 in the right hemisphere (images in Fig. 3A and Fig. S1 were mirrored for consistency with cited references). Virus injections were performed at 3–5 sites into the binocular region of V1 and ~0.5–1 µm more lateral (corresponding to the location of areas LM and RL), using AAV2/1.Syn.mRuby2.GSG.P2A.GCaMP6s.WPRE.SV40 (Rose et al. 2016), diluted to reach a final titer of ~1.5e-13 genome copies/ml. The virus solution was injected using glass pipettes (tip diameter, 10–40 µm) and a pressure micro-injection system, at 200–450 µm below the cortical surface (100–150 nl/injection, 20 nl/min pressure injected at 0.25 Hz). Following injections, the craniotomy was sealed flush with the brain surface using a glass cover slip (4 or 5 mm diameter) and cyanoacrylate glue (Histoacryl), avoiding glue contacting the brain. A custom machined aluminum head-plate was attached to the skull using dental cement to allow head-fixation during imaging. Expression of the transgene was allowed for 2.5–3 weeks before imaging.

### Intrinsic signal imaging

Intrinsic signal imaging was used to localize areas V1, LM, and RL. Imaging was performed 2-4 weeks after cranial window implantation, under anesthesia by intraperitoneal injection of a mixture of Fentanyl (0.030 mg/kg), Midazolam (3.0 mg/kg), and Medetomidine (0.30 mg/kg). The surface of the brain was illuminated through the cranial window with red light from two sides using a 735-nm LED (bandpass filtered at 700/40 nm). Images were collected through a 4× air objective (NA 0.28, Nikon) using a CCD camera (Teledyne Dalsa Xcelera-LVDS PX4,; 12 bit; 512×512 pixels; 15Hz; spatial binning, 3×3 pixels; temporal binning, 3 frames). The imaging plane was set 400–500 µm below the cortical surface. In addition, an image of the cortical surface was acquired using green light from a 530-nm LED to visualize the blood vessel pattern, which was used as a reference to target two-photon imaging. Acquisition and analysis software were custom written in Matlab.

### In vivo two-photon imaging

Two-photon imaging was performed 3–23 weeks (median, 4.7 weeks, IQR 4.3-5.9) after cranial window implantation under anesthesia. Mice were initially anesthetized by intraperitoneal injection of a mixture of Fentanyl (0.030 mg/kg), Midazolam (3.0 mg/kg), and Medetomidine (0.30 mg/kg). Additional anesthetic mixture (25% of the induction dose) was injected subcutaneously 60 min after the initial injection and then every 30-40 min to maintain anesthesia. Mice were placed on a heated blanket to ensure thermal homeostasis and fixed through the head-plate under the microscope. Images were acquired using a custom-built two-photon microscope equipped with an 8 kHz resonant galvanometer scanner operated in bidirectional mode, resulting in frame rates of 17.6 Hz at an image resolution of 750×900 pixels (330×420 µm). The illumination source was a Ti:Sapphire laser with a DeepSee pre-chirp unit (Spectra Physics MaiTai eHP, <100 fs pulse width, 80 MHz repetition rate), set to an excitation wavelength of 940 nm. Laser power was modulated with a half-wave plate combined with a polarizing beam splitter cube, and was between 8–25 mW as measured after the objective (16×, 0.8 NA, Nikon) with a photodiode. A mechanical blanker was positioned in the focal plane between the scan and tube lenses to block the laser beam at the scan turnaround points. Photons collected from the objective passed through a beam splitter (FF560 dichroic) and were directed onto two separate photomultiplier tubes (PMT, Hamamatsu R6357) with a green (525/50-25 nm) and red (607/70-25 nm) band pass emission filter. Data were acquired with a high-speed digitizer (NI-5761, National Instruments, 500 MHz) in combination with a field-programmable gate array (FPGA) to bin the PMT signal into pixels. In some cases, mice were repeatedly imaged to increase yield. When a given visual area was targeted for a second time in the same animal, the new imaging plane was acquired at least 20 µm below or above the previously imaged plane, which could be readily reidentified using the structural marker mRuby2, thereby ensuring that there was no double sampling of cells. For awake imaging the animal was head-fixed on top of an air suspended Styrofoam ball (diameter 20 cm), allowing the mouse to run freely during stimulus presentation and data acquisition (Dombeck et al. 2007).

### Monitoring eye position

During two-photon imaging, both eyes were continuously imaged with an infrared video camera (The Imaging Source, frame rate 30 Hz). Pupil position and diameter were monitored online using custom-written software (LabVIEW, National Instruments) based on Sakatani and Isa (2007). Analysis of pupil position was also performed post hoc to test whether either eye had changed position over the course of the experiment. Approximately 10% of the experiments were discarded owing to eye drifts.

### Visual stimulation

#### Retinotopic mapping with intrinsic signal imaging

To locate the binocular region of V1 for cranial window positioning, retinotopic mapping was performed using small patches of drifting gratings (size 20–30 deg) presented at 8 to 15 different locations on a gray background (eight consecutive directions in a pseudorandom sequence; spatial frequency (SF), 0.04 cpd; temporal frequency (TF), 2 Hz; duration of each stimulus patch, 6 sec; inter-stimulus interval, 3 sec; number of stimulus trials, 3–4). The stimulus monitor (27 inches, Dell SE2717H, gamma corrected, refresh rate 60 Hz, spatial resolution 1920×1080 pixels) was obliquely placed in the left (contralateral) visual hemifield, approximately 30 deg from the mouse’s midline at a distance of 12 cm from the left eye, covering approximately from –20 to +100 deg in azimuth and from –40 to +40 deg in elevation.

#### Mapping higher visual areas with intrinsic signal imaging

To locate higher areas of mouse visual cortex, retinotopic mapping employing a periodically drifting bar was used for both cardinal axis (Kalatsky and Stryker 2003; Marshel et al. 2011). The bar consisted of a reversing checkerboard pattern (SF, 0.04 cpd; TF, 2 Hz) and was periodically swept over a gray background at 18–20 deg/sec, 30–45 times for each direction. Bar width was 20 deg, and spherical correction was applied to stimulate in spherical visual coordinates using a flat monitor. The stimulus monitor (27 inches, Dell SE2717H, gamma corrected, refresh rate 60 Hz, spatial resolution 1920×1080 pixels) was obliquely placed in the left (contralateral) visual hemifield, at an angle of approximately 30 deg from the mouse’s midline and a distance of 12 cm from the left eye, covering approximately from –20 to +100 deg in azimuth and from –40 to +40 deg in elevation.

#### Dichoptic stimulation

Eye shutter glasses (3D Vision 2, Nvidia) were used for independent stimulus presentation to each eye. The glasses consisted of a pair of liquid crystal shutters, one for each eye, that rapidly (60 Hz) alternated their electro-optical state – i.e. either occluded or transparent to light. In one frame sequence, the left eye shutter is occluded while the right eye shutter is transparent, and vice versa for the next frame, with alternations synchronized to the monitor refresh rate (120 Hz). Synchrony between the shutter glasses and the monitor was accomplished with an infra-red wireless emitter. The display monitor (Acer GN246HL, 24 inches, 120 Hz refresh rate, spatial resolution 1600×900 pixels) was placed in front of the mouse at a distance of 13 cm from the eyes (luminance measured through the transparent shutter: white, 21.6 cd/m^2^; black 0.05 cd/m^2^). To reduce light contamination of two-photon images from visual stimulation, the microscope objective was shielded using black tape. Visual stimulation and shutter control were based on custom-written code for Matlab (MathWorks) using the Psychophysics Toolbox (Brainard 1997; Pelli 1997; Kleiner et al. 2007), running on a Dell PC (Precision T7500) under Windows 10 and equipped with a Nvidia Quadro K4000 graphics card.

#### Dichoptic drifting gratings

Drifting vertical gratings were dichoptically presented to both eyes, at varying interocular disparities. Different interocular grating disparities were generated by varying the initial phase (position) of the grating presented to one eye relative to the phase of the grating presented to the other eye across the full grating cycle. Twelve equally spaced phase disparities (−150–180 deg phase, spacing 30 deg phase) were used. For each stimulus, drift direction (leftward or rightward), TF (2 Hz), and SF (0.01 cpd) were kept constant across eyes. Each stimulus was displayed at 70% contrast for 2 sec (4 grating cycles, randomized initial spatial phase) preceded by an inter-stimulus interval of 2 sec with a blank (gray) screen with the same mean luminance as during the stimulus period. Gratings were presented in pseudorandomized sequence across disparities and drifting directions, with 5-6 trials for each stimulus condition.

#### Random dot stereograms

Random dot stereograms (RDS) consisted of a pattern of random dots, presented to both eyes in a dichoptic fashion. Between the left and the right eye stimulus patterns, a spatial offset along the horizontal axis was introduced to generate interocular disparities. A total of 23 different RDS conditions were presented, covering a range of disparities between –31.3 deg and +31.3 deg. The different RDS conditions were obtained by dividing the entire range of disparities into 23 nonoverlapping bins (bin width 2.6 deg) and assigning each bin to one RDS condition (e.g. [–1.3 +1.3], [+1.3 +3.9], etc.). Each RDS stimulus was presented for 5 sec, during which a new random pattern of dots was displayed every 0.15 sec. In each pattern, all dots had the same interocular disparity, randomly chosen within the 2.6 deg bin of that particular RDS condition. The dots (diameter 12 deg) were bright (brightness 77%) against a gray background with an overall density of 25%. Each RDS condition was presented for 9–10 stimulus trials, with individual trials separated by an inter-stimulus interval of 2 sec. RDS stimuli were presented in pseudorandomized sequence, interleaved with dichoptic gratings as part of the same stimulation sequence.

#### Receptive field mapping

Receptive field (RF) elevation for individual neurons was determined using horizontal bars of drifting gratings displayed at 11 different vertical locations on a gray background (four consecutive cardinal directions in non-random sequence; SF, 0.03 cpd; TF, 2 Hz; contrast, 60%; duration of each stimulus bar, 4 sec; inter-stimulus interval, 1 sec; number of stimulus trials, 6). The bar width was 14.6 deg and adjacent locations were 7.3 deg apart (50% bar overlap), covering approximately from –45 to 42 deg in elevation in total. Bar stimuli were presented to each eye separately using eye shutter glasses, in pseudorandomized sequence.

### Data analysis

All data analyses were performed using custom-written Matlab code (MathWorks).

#### Image analysis

Imaging data were processed in three steps: (1) image registration, (2) selection of regions of interest (ROIs), (3) extraction of calcium fluorescence time courses. (1) Motion artifacts, mainly consisting of small, slow drifts in brain position that occurred during imaging, were corrected by rigid translational registration based on 2D cross-correlation and applied to down-sampled fast Fourier transforms of all frames, using the average of the initial 200 frames of the recording as a reference. (2) ROIs were selected by manually drawing circular shapes around cell somas, which were morphologically identified by inspecting the average of all registered frames of the recording, combined with examination of activity maps. Regions with overlapping somas were excluded from the analysis. (3) Since the signal from any somatic ROI might be contaminated by out-of-focus fluorescence from surrounding neuropil and other cells, neuropil contamination was corrected by generating a peri-somatic neuropil ROI for each soma ROI, consisting of an annular region extending from 3 µm (7 pixels) to 13 µm from the border of the somatic ROI. Pixels belonging to other somatic ROIs were excluded from neuropil ROIs. The raw fluorescence time course of each cell was extracted by averaging all pixels within the somatic ROI (F_cell_raw_). Similarly, the fluorescence time course from the annular neuropil ROI (F_neuropil_) was extracted. The true fluorescence time course of a cell was estimated as F_cell_corrected_ = F_cell_raw_ − *r* × F_neuropil_, with a contamination factor *r* set to 0.7 (Kerlin et al. 2010; Chen et al. 2013). For a subset of experiments, imaging data were processed using the Suite2P toolbox in Matlab (Pachitariu et al. 2016), which entailed image registration, segmentation of ROIs, and extraction of calcium fluorescence time courses. All automatically segmented ROIs were manually curated to exclude ROIs not corresponding to somata, based on morphology and by inspecting the average of all registered frames of the recording. After neuropil correction, the fluorescence signals were filtered with the Savitsky-Golay method (second order polynomial, 10 data points, 0.5 sec). Relative changes in fluorescence signals (ΔF/F_0_) were calculated, for each stimulus trial independently, as (F *−* F_0_)/F_0_, where F_0_ was the average over a baseline period of 1 sec immediately before onset of the visual stimulus.

#### Retinotopic mapping of V1

Images were high-pass-filtered to calculate blank-corrected image averages for each stimulus condition, and thresholded (image background mean + 3 × standard deviation). The signal in response to visual stimulation (23 frames during presentation of a given visual stimulus) was referenced against the mean of 15 baseline frames (prior to stimulus presentation), resulting in a percentage signal decrease for each pixel. A map of retinotopy was generated by assigning a color to each pixel based on the stimulus location that evoked the strongest response and encoding the response strength by pixel intensity.

#### Retinotopic mapping of higher visual areas

Retinotopic maps for azimuth and elevation were generated using the temporal phase method introduced by Kalatsky and Stryker (Kalatsky and Stryker 2003) on images obtained with intrinsic signal imaging. Briefly, pixels with a similar temporal phase in response to the vertical or horizontal periodic bar encode, respectively, iso-azimuth or iso-elevation coordinates in the visual field. To compute maps, first, the time course of each pixel was high-pass filtered using a moving average (with a time window equaling the duration of the moving bar cycle, ~10 sec) to remove slow artifactual changes in reflected light intensity not evoked by visual stimulation. Next, a Fourier transform was computed to extract the phase and the power of the frequency component at the bar drifting frequency. The phase indicates the location of the bar driving the response of a pixel, and the power indicates the strength of its response. To compute maps of absolute retinotopy, the response time to the bar drifting in one direction was subtracted from the response time to the opposite drift direction (). From these maps of absolute retinotopy, equally spaced iso-azimuth and iso-elevation contour lines were extracted, color-coded for visual field location, and overlaid on top of the image of the blood vessel pattern. The boundary between V1 on the medial side and areas LM, AL, and RL on the lateral side was identified by a reversal at the vertical meridian, as indicated by the longer axis of the elliptically shaped contour on the vertical meridian. The boundaries between LM and AL, and between AL and RL were identified as a reversal near the horizontal meridian (; Marshel et al. 2011; Garrett et al. 2014). The binocular regions of areas V1, LM, and RL were then specifically targeted for two-photon imaging, by using the blood vessels as landmarks, which could be reliably recognized in the two-photon images.

#### Responsive cells

Cells were defined visually responsive when ΔF_peak_/F_0_ > 4 × *σ*_baseline_ in at least 50% of the trials of the same stimulus condition, where ΔF_peak_ is the peak ΔF/F_0_ during the stimulus period of each trial, and *σ*_baseline_ is the standard deviation calculated across the F_0_ of all stimulus trials and conditions of the recording. For grating stimuli, the mean ΔF/F_0_ over the entire stimulus interval (2 sec) of each trial was calculated. For RDS stimuli, the mean ΔF/F_0_ of each trial was calculated over a time window of 1 sec centered around ΔF_peak_. When plotting the disparity preference averaged across neurons of each imaging plane (Fig. 2A,C), only imaging planes with at least 10 disparity-tuned neurons were used for averaging.

#### Disparity selectivity index

For each cell responsive to dichoptic gratings, a disparity selectivity index (DI) was calculated, given by the normalized length of the mean response vector across the twelve phase disparities of the drifting direction that elicited the stronger activation (Scholl et al. 2013; 2015):

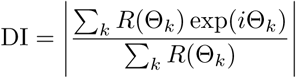

where *R*(Θ_*k*_) is the mean ΔF/F response to the interocular phase disparity Θ_*k*_. Cells were defined disparity-tuned if DI>0.3. Using more stringent criteria for defining responsive (ΔF_peak_/F_0_ > 8 × *σ_baseline_*) did not result in a significant change in the DI distribution for each area (data not shown), indicating that signal-to-noise issues did not affect the measurement of disparity selectivity. Note that the calculation of DI is based on a circular metric. As such, DI could be computed only for responses to dichoptic gratings, but not for responses to RDS, which are not circular. Cells were defined disparity-tuned to RDS when at least 50% of the tuning curve variance (*R*^2^) could be accounted for by the model fit (see below).

#### Disparity tuning curve fit

Disparity tuning curves obtained with either dichoptic gratings or RDS were fitted with an asymmetric Gaussian function using single trial responses, as follows:

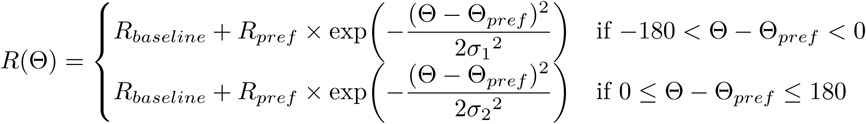

where *R*_*baseline*_ is the baseline response, *R*_*pref*_ is the response to the preferred disparity, *σ*_1_ and *σ*_2_ are the tuning width parameters for the left and right sides, respectively. The tuning width as plotted in Fig. 1F was calculated as *σ*_1_ + *σ*_2_. Disparity preferences of responsive cells for either dichoptic gratings or RDS were given by the fit parameter *Θ*_*pref*_ when at least 50% of the tuning curve variance (*R*^2^) could be accounted for by the model fit.

#### Receptive field elevation

To determine RF elevation, only the eye that elicited the strongest activation of a neuron was used. A one-dimensional RF was obtained by fitting the responses of each cell as a function of the stimulus positions with a Gaussian model, and the position in elevation corresponding to the RF peak was taken when at least 50% of the tuning curve variance (*R*^2^) could be accounted for by the model fit.

### Statistics

Statistical analyses were performed using Matlab (MathWorks). Sample sizes were not estimated in advance. No randomization or blinding was performed during experiments or data analysis. Data are reported as mean with standard error of the mean (mean ± SEM), or as median with interquartile range (median with 25^th^ and 75^th^ percentiles), as reported in the Figures and Figure legends. For calculations across imaging planes, only imaging planes with at least 10 responsive cells (Fig. 1C) or responsive and disparity-tuned cells (Fig. 1F, Fig. 2B,D, Fig. 3) were considered. Data groups were tested for normality using the Shapiro-Wilk test in combination with a skewness test and visual assessment (Ghasemi and Zahediasl 2012). Quantifications for data obtained with grating phase disparities were performed taking into account their circularity (Berens 2009). Before calculating medians for circular data, the Rayleigh test and the Omnibus test were used to verify deviations from circular uniformity. Comparisons between data groups where made using the appropriate tests (Kolmogorov-Smirnov test, Watson-Williams test, one-way ANOVA, Kruskal-Wallis test). For multiple comparisons, Bonferroni correction was used. Significance levels in Fig. 3E,H were determined by permutation tests, by randomly shuffling labels between the two data groups (V1 and RL) and computing the difference between group means (or circular means) for each bin in each permutation; p values were computed as the proportion of permutations (n=10000 permutations) with more extreme mean differences than the observed ones. All tests were two-sided. Correlation coefficients were calculated as Spearman’s correlation coefficient or as correlation coefficient between a circular and linear variable. The statistical significances are reported in the Figures, with asterisks denoting significance values as follows: * p < 0.05, ** p < 0.01, *** p < 0.001.

### Data and code availability

Data and analysis code used in this study are stored and curated at the Max Planck Computing and Data Facility Garching (Munich, Germany) and are available from the corresponding authors upon reasonable request.

### Supplementary Figures

**Figure S1.**
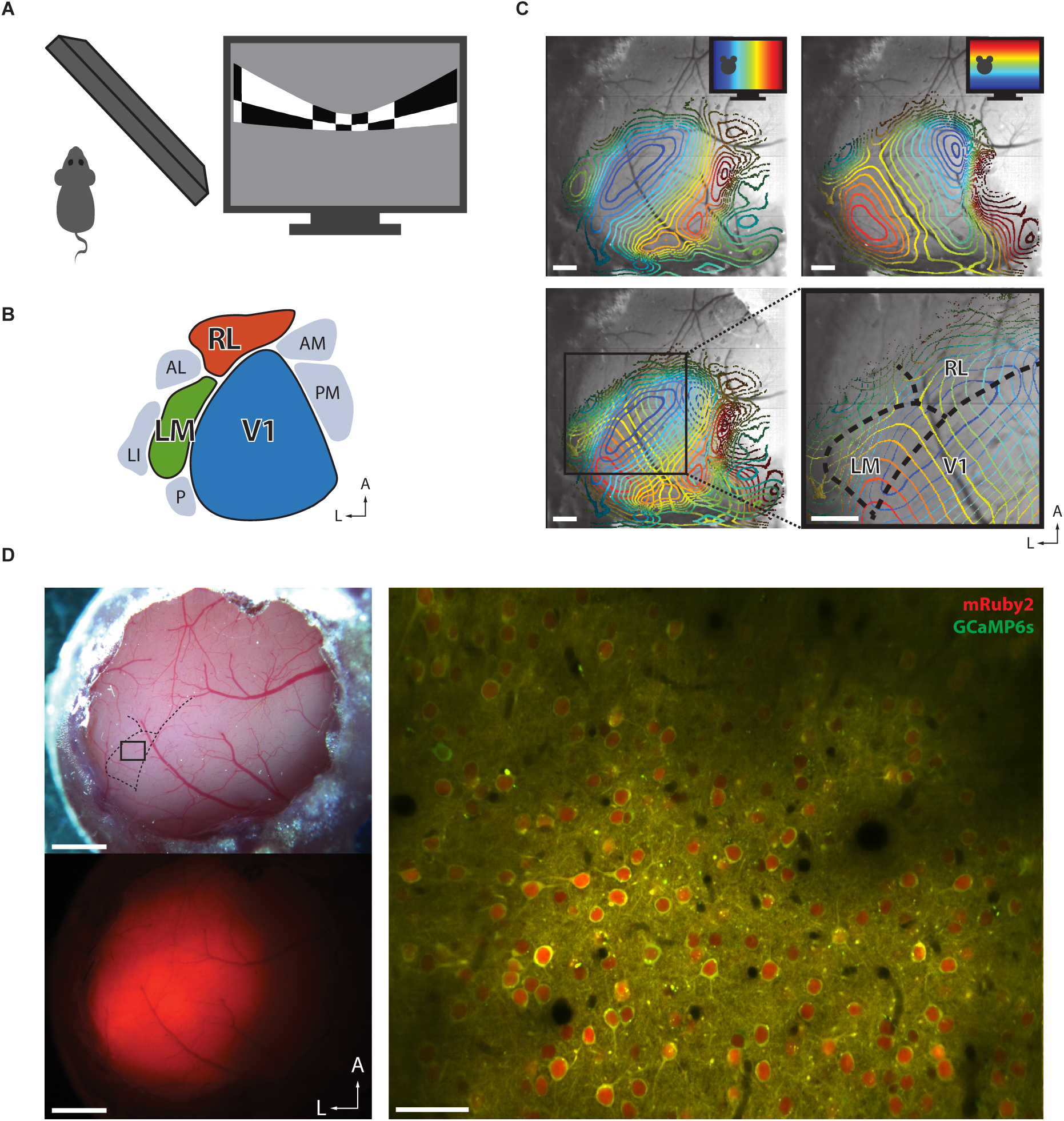
Identification and targeting of areas V1, LM, and RL for two-photon calcium imaging. (A) Schematic of stimulus presentation for mapping the retinotopic organization of mouse visual cortex areas. Left, top view; right, periodic bar stimulus displayed with spherical correction. (B) Schematic of the location of V1 and several higher-order areas of mouse visual cortex in the left hemisphere. The color code for areas V1 (blue), LM (green), and RL (orange) is used throughout the Figures. (C) Retinotopic maps from an example mouse. Contour plots of retinotopy are overlaid with an image of the brain surface. Contour lines depict equally spaced, iso-elevation and iso-azimuth lines as indicated by the color code. Top left, contour plot for azimuth; top right, contour plot for elevation. Bottom left, overlay of azimuth and elevation contours. Bottom right, enlarged view of cortical areas V1, LM, and RL. The boundaries between these areas (dashed black lines) can be reliably delineated. Scale bars, 500 µm. (D) Two-photon imaging using the calcium indicator GCaMP6s co-expressed with the structural marker mRuby2. Top left, image of a cranial window 5 weeks after implantation. Bottom left, epifluorescence image showing the expression bolus, with fluorescence signal from mRuby2. Right, example two-photon imaging plane acquired 180 µm below the cortical surface in area LM. The image shows a mean-intensity projection (19000 frames, shift corrected) with fluorescence signal from GCaMP6s (green) and mRuby2 (red). The cortical location of this imaging plane is indicated in the top left panel. Scale bars: left panels, 1 mm; right panel, 50 µm.

**Figure S2.**
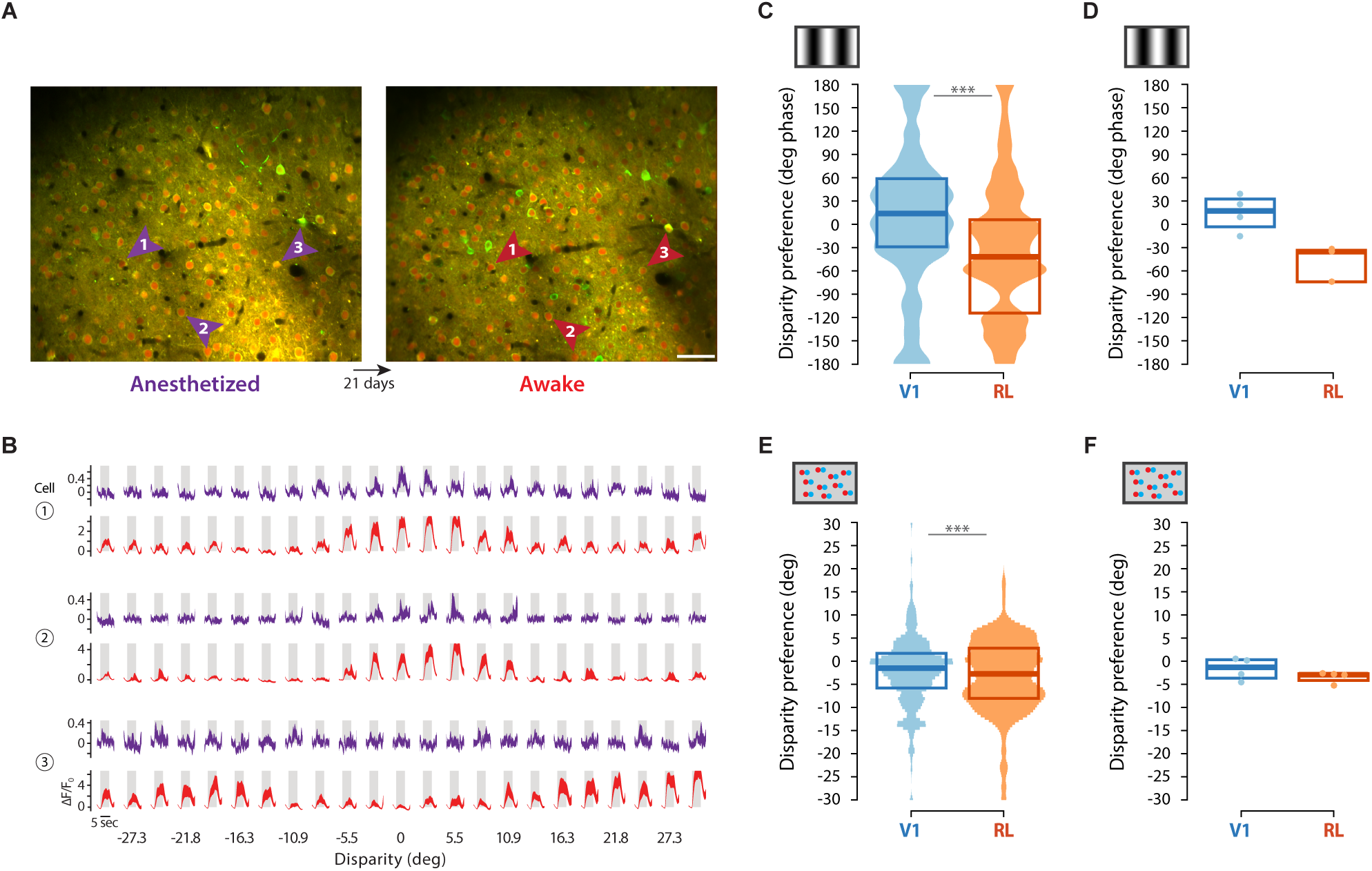
Imaging in awake mice reveals strong disparity-tuned responses to RDS and near disparity tuning in RL. (A) Re-finding of the same neurons in two imaging sessions. Left, frame-averaged two-photon image of an imaging plane recorded in V1 in the anesthetized animal. Right, same imaging plane, acquired 21 days later in the awake animal. Scale bar, 50 µm. (B) Comparison of visually-evoked responses between the anesthetized and the awake state. Calcium traces (ΔF/F_0_) in response to RDS stimuli of three example neurons (indicated in (a)) measured in the anesthetized (purple) and awake (red) animal. Shaded regions represents ± SEM calculated across stimulus trials (10 repeats). (C) Disparity preference measured with gratings, averaged across all disparity-tuned neurons in areas V1 and RL of awake mice. Box plots indicate median and interquartile range calculated across neurons. Shaded areas show the mirrored and normalized circular probability density estimates, kernel width 10 deg (V1: n=135 cells, 4 imaging planes, 3 mice; RL: n=107 cells, 4 imaging planes, 2 mice; Watson-Williams test, F(1,240)=21.167, p < 6.8306e-06). (D) Disparity preference measured with gratings, averaged across neurons in each imaging plane. Box plots indicate median and interquartile range calculated across imaging planes. Individual data points show the mean disparity preference calculated across neurons in each imaging plane (V1: n=4 planes, 3 mice; RL: n=3 planes, 2 mice; statistical tests were not performed owing to the small sample size). (E) Disparity preference measured with RDS, averaged across all disparity-tuned neurons in each area. Box plots indicate median and interquartile range calculated across neurons. Shaded areas show the mirrored and normalized probability density estimates, kernel width 3 deg (V1: n=356 cells, 4 imaging planes, 3 mice; RL: n=262 cells, 4 imaging planes, 2 mice; Kolmogorov-Smirnov test: p = 2.5738e-03). (F) Disparity preference measured with RDS, averaged across neurons in each imaging plane. Box plots indicate median and interquartile range calculated across imaging planes. Individual data points show the mean disparity preference calculated across neurons in each imaging plane (V1: n=4 planes, 3 mice; RL: n=4 planes, 2 mice; statistical tests were not performed owing to the small sample size).

**Figure S3.**
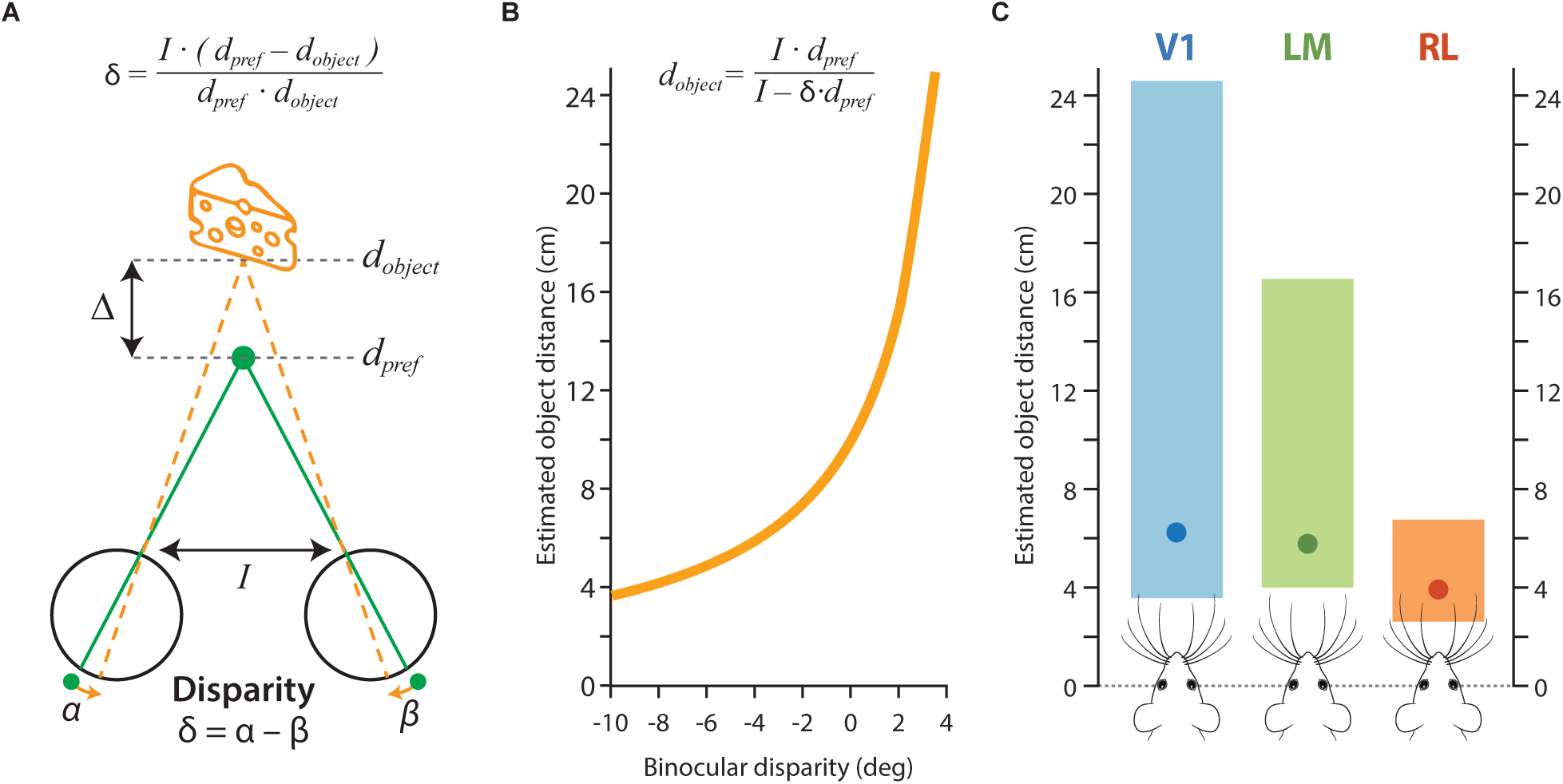
Area RL is estimated to cover a narrow range of distances, very close to the mouse. (A) Stereo-geometry of the mouse visual system (adapted, with permission, from Scholl et al. 2013). Absolute object distance can be calculated from the binocular disparity caused by the object, following the equation given by Scholl et al. (2013), using the values of preferred binocular viewing distance (*d*_*pref*_ = 10 cm) and interocular distance (*I* = 1.0 cm), and with object depth Δ defined as *d_pref_ − d_object_*. The preferred binocular viewing distance is estimated to be 10 cm, based on the fact that the refractive error of mouse eye of +10.0 diopters enable optimal focusing of objects at a distance of 10 cm (Cera et al. 2006). (B) Relationship between binocular disparity and absolute object distance, derived from the equation in (a). Note that using different values of preferred binocular viewing distance *d*_*pref*_, e.g. 5 cm or 20 cm, has only little effect on the estimated object distances (data not shown). (C) Range of distances encoded by disparity-tuned neurons in areas V1, LM, and RL. The range of distances is estimated by using the equation in (a) to transform the interquartile ranges of preferred disparities as shown in Fig. 2D (V1: 3.6–24.9 cm, LM: 4.0–16.7 cm, RL: 2.6–6.8 cm). Dots are the estimated distances corresponding to the median disparity preference across neurons (Fig. 2D; V1: 6.3 cm, LM: 5.8 cm, RL: 3.9 cm). In the mouse schematics at the bottom, the eyes are aligned at zero (dotted line), with the interocular distance (1 cm) and whiskers in scale with the y-axis.

